# Non-invasive stimulation with Temporal Interference: Optimization of the electric field deep in the brain with the use of a genetic algorithm

**DOI:** 10.1101/2022.02.10.477970

**Authors:** D. Stoupis, T. Samaras

## Abstract

**Objective:** Since the introduction of transcranial temporal interference stimulation (tTIS), there has been an ever-growing interest in this novel method, as it theoretically allows non-invasive stimulation of deep brain target regions. To date, attempts have been made to optimize the electrode montages and injected current to achieve personalized area targeting using two electrode pairs. Most of these methods use exhaustive search to find the best match, but faster and, at the same time, reliable solutions are required. In this study, the aforementioned stimulation settings were optimized using a genetic algorithm.

**Methods:** Simulations were performed on head models from the Population Head Model (PHM) repository. First, each model was fitted with an electrode array based on the 10-10 international EEG electrode placement system. Following electrode placement, the models were meshed and solved for all single-pair electrode combinations using an electrode on the left mastoid as a reference (ground). At the optimization stage, different electrode pairs and injection currents were tested using a genetic algorithm to obtain the optimal combination for each model.

**Results:** Greater focality was achieved with optimization, specifically for the right hippocampus (region of interest), with a significant decrease in the surrounding electric field intensity. In the non-optimized case, the brain volumes stimulated above 0.2V/m were 99.9% in the region of interest (ROI), and 76.4% in the rest of the gray matter. In contrast, the stimulated volumes were 91.4% and 29.6% for the best optimization case. Additionally, the maximum electric field intensity inside the region of interest was consistently higher than that outside the ROI for all optimized cases.

**Significance:** Given that the accomplishment of a globally optimal solution requires a brute-force approach, the use of a genetic algorithm can significantly decrease the optimization time and provide usable results. Although the method is promising and the indications are becoming more robust, such that optimal solutions can be achieved at the individual level, there are limitations on what can be achieved using only two electrode pairs.

## 1. Introduction

The anatomical and functional complexity of the human brain makes the electrical stimulation of neurons located in specific areas a challenging task. Invasive stimulation is an obvious solution in some cases; however, it poses considerable risk to the patient. Non-invasive methods have been proposed to stimulate specific areas deep in the brain to minimize the risk induced by invasive methods.

The most common non-invasive brain stimulation techniques include direct electric field induction methods, such as transcranial direct current stimulation (tDCS), transcranial alternating current stimulation (tACS), and magnetically induced electric field methods, namely, transcranial magnetic stimulation (TMS). The efficacy of such methods in specific psychiatric conditions ranges from promising to indisputable [1]–[4]. The main problem with all of the above is the depth at which stimulation is effective. Various targets, such as the thalamus or hippocampus, lie deep within the brain, and new ways to stimulate these areas non-invasively are yet to be fully developed. Such a method is the transcranial temporal interference stimulation (tTIS), initially proposed by Grossman et al. [5]. This technique takes advantage of the spatial electromagnetic wave interference resulting from stimulation frequencies in the 1 - 100 kHz range. As neurons do not respond to higher than 1 kHz frequencies [6], the effect of the individual electrode high-frequency currents is minimal; thus, only the modulation envelope of the temporal interference stimulates the neurons (for a small frequency difference). The mechanisms by which this method works are still under active research, with the most recent developments described by Mirzakhalili et al. [7] and Esmaeilpour et al. [8].

There has been an increasing interest in the tTIS method, which, as Grossman et al. [5] demonstrated, works well in murine subjects. More specifically, current research on this method seeks to optimize the electrode combinations and currents [9], [10]. Recognizing this problem in the present work, we aim to perform a study on the optimization effectiveness and present a way of optimizing the effects of tTIS at any head model generated from MRI data. The main deep brain areas investigated in this study were the hippocampus and the thalamus. The first is the optimization target, and the second serves as a control area for focality and effectiveness. The hippocampus on the right hemisphere was chosen as the optimization target since we can obtain results comparable to those of other studies [9], [10]. The thalamus, on the other hand, covers a significantly larger area than the hippocampus and lies in its close vicinity; therefore, it can be used as a control area for optimization, expecting to observe a significant decrease in the maximum electric field in the thalamic area in the case of an acceptably optimized electric field in the unilateral hippocampal area.

Two different optimization measures were used in this study. The first is the ratio of the electric field in the region of interest to the electric field in the rest of the brain, called ROI/Rest. The second measure was the maximum electric field in the ROI and in the control region, which in this study were the right hippocampus and thalamus, respectively. The ROI/Rest measure is also the optimization objective function, and it is presented in more detail in the Materials and Methods section.

## 2. Materials and Methods

### 2.1. Simulation of tTIS electric field

For the tTIS stimulation technique, only the electric field is of interest. In the case of low-frequency current injected into the electrodes, the domain can be considered ohmic quasi-static [11] without any significant displacement current [12]. This approximation simplifies the equations and, assuming there are no internal current sources, the Laplace equation in the following form is the one solved in the finite element (FE) computational domain:

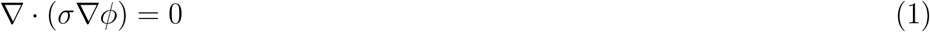

with *σ* being the electrical conductivity (S/m) of each tissue.

The conductivity value may vary depending on the frequency of the field or individual characteristics, such as age, gender, and pathology. Conductivity values suggested by McCann et al. [13] were used this study. Equation (1) was integrated over the entire volume, to calculate electric potential at each node. All finite element model (FEM) calculations were executed with SfePy [14] using the Portable, Extensible Toolkit for Scientific Computation (PETSc) package [15]–[17], with the conjugate gradient (CG) solver and an algebraic multigrid (BoomerAMG) matrix preconditioner from the High-Performance Preconditioners (HYPRE) [18]. Dirichlet boundary conditions were used on the electrodes, setting the active electrode to 1V and the return electrode to −1V. After the solution, the electric field values were scaled by the ratio of the desired electric current with the current passing through the electrode-skin interface surface.

Using two pairs of electrodes (Figure 1), with each pair having a slightly different frequency, an interference pattern can be generated in the conducting medium, whose envelope oscillates at the difference between the two frequencies. Depending on the nature of the medium, the pattern can vary, and as illustrated by Grossman et al. [5], it can easily be calculated when a uniform medium is used.

**Figure 1.**
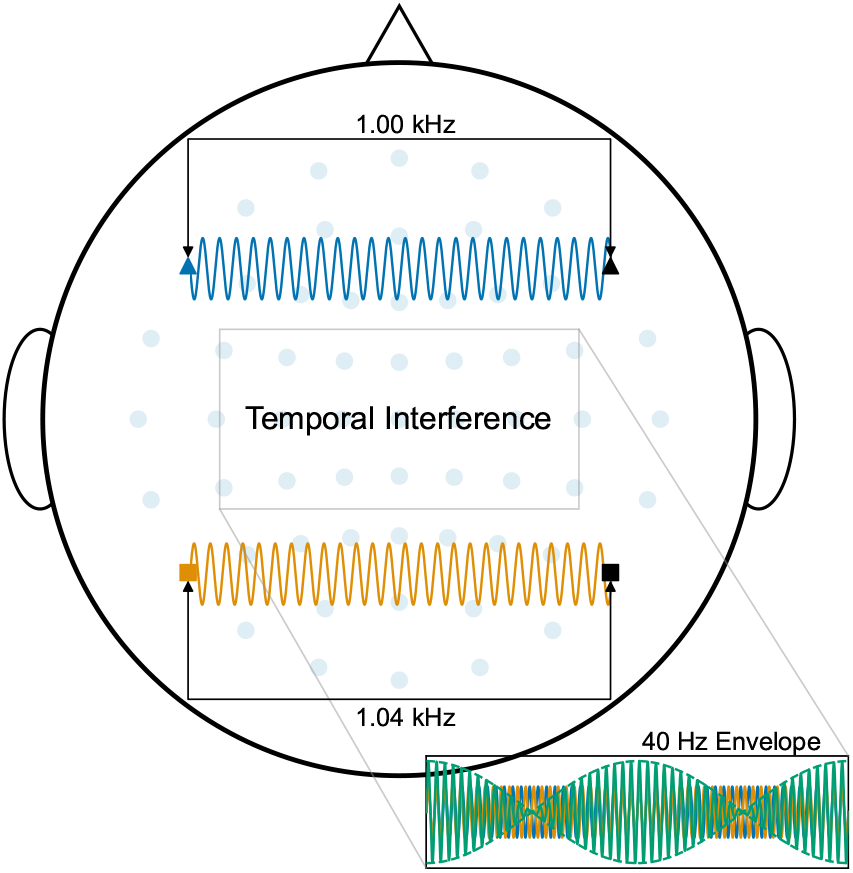
Demonstration of the Temporal Interference Stimulation method. Electrodes annotated as Δ and □ are two distinct pairs. The black part of the electrode pairs resembles the ground.

Since the modulation occurs in 3D space, there will be different patterns in the *x*, *y*, and *z* directions. According to Grossman et al. [5, page 20], the envelope amplitude of the amplitude modulation (AM) of the electric field produced by the temporal interference at any location 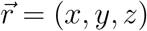, is calculated as

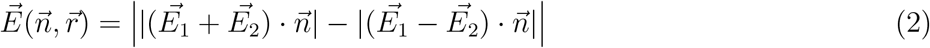

where 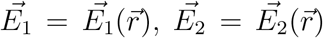 are the electric fields coming from the two electrode pairs and 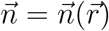 is the unit vector in the direction of interest.

The maximum amplitude of the modulation envelope at a specific location is of primary interest in this study because the modulation varies with time between zero and the maximum. Starting from Equation (2), the equations for calculating the maximum modulation amplitude across all directions at a specific location 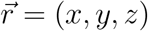 are [5, page 20]:

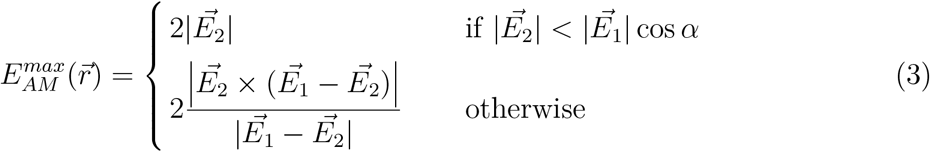

where *α* is the angle between 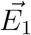 and 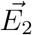, and Equation (3) holds only if *α* < 90°.

When *α* ≥ 90°, the sign of one of the two fields can be flipped (it must be done consistently) since reaching peak field strength at different time points across different areas violates the rule of less than 90°. This change is possible, considering that Equation (3) calculates the maximum effect over one oscillation.

### 2.2. Realistic Head Models

Realistic head models are required to generate accurate results for the field distribution, and such models can be generated from MRI scans of individuals. Obtaining meaningful results requires the use of several different head models. The Population Head Models (PHM) repository is a valuable resource with 50 head models from healthy individuals aged 22-35 years, with an anatomically realistic grey matter structure. Of these 50 models, 38 were used in this study, and their IDs are listed in Table 1. The omitted models had insolvable self-intersections in their generated surface meshes, rendering numerical calculations impossible.

**Table 1.**
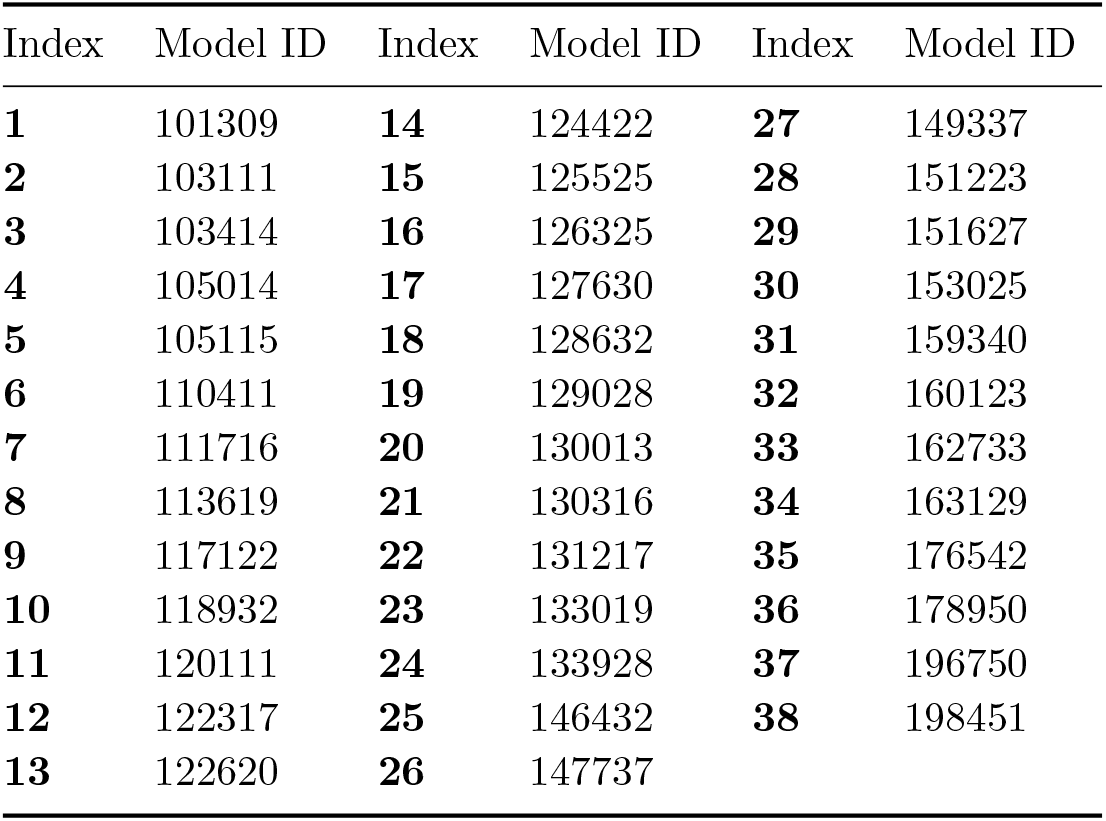
Index assignment to each model, based on the ID from the PHM repository. All the figures of this work will follow this convention, unless a different notation is explicitly used.

| Index | Model ID | Index | Model ID | Index | Model ID |
| --- | --- | --- | --- | --- | --- |
| **1** | 101309 | **14** | 124422 | **27** | 149337 |
| **2** | 103111 | **15** | 125525 | **28** | 151223 |
| **3** | 103414 | **16** | 126325 | **29** | 151627 |
| **4** | 105014 | **17** | 127630 | **30** | 153025 |
| **5** | 105115 | **18** | 128632 | **31** | 159340 |
| **6** | 110411 | **19** | 129028 | **32** | 160123 |
| **7** | 111716 | **20** | 130013 | **33** | 162733 |
| **8** | 113619 | **21** | 130316 | **34** | 163129 |
| **9** | 117122 | **22** | 131217 | **35** | 176542 |
| **10** | 118932 | **23** | 133019 | **36** | 178950 |
| **11** | 120111 | **24** | 133928 | **37** | 196750 |
| **12** | 122317 | **25** | 146432 | **38** | 198451 |
| **13** | 122620 | **26** | 147737 |  |  |

Each model comprises eight surface meshes: skin, skull, cerebrospinal fluid (CSF), gray matter, white matter, cerebellum, and ventricles. The tetrahedral mesh of each model was generated using TetGen [19]. The meshing parameters for TetGen were chosen based on the study by Tran et al. [20], considering the computational resources required. The final meshes were produced with *q* = 1.4 (the tetrahedral radius-edge ratio) and *v_max_* = 8 (the maximum individual tetrahedron volume).

All models were fitted with electrodes based on the international 10-10 EEG electrode placement system. The four required fiducials, nasion, inion, and right and left preauricular, were individually identified for each model, and Mesh2EEG [21], [22] was used to calculate the positions of the electrodes on the skin of each model, yielding a total of 61 electrodes. An additional electrode was placed on the left mastoid, serving as a reference electrode (P9 in the international 10-10 system).

### 2.3. Automatic Anatomical Labeling (AAL) Atlas Registration

Atlas labeling is required to perform meaningful comparisons between different models, since geometry varies significantly. To efficiently perform region analysis, the solved models were converted to the Neuroimaging Informatics Technology Initiative (NIfTI) format and then normalized to the Montreal Neurological Institute (MNI) space using the Statistic Parametric Modelling (SPM) library [23] in MATLAB. Each model had a resolution of 1 mm per voxel, and after normalization, the individual brain areas were identified using the AAL atlas v3.1 [24].

### 2.4. Optimizing Electrode Montages and Currents

Electrode position and current intensity optimization have been well studied in tDCS [25], [26], although the problem in tTIS is not convex. Owing to the nonlinearity and non-convex nature of the tTIS method, exhaustive search methods [9], [10] are usually employed to achieve personalized optimization. We used an evolutionary algorithm to determine the optimal electrode and current combination for the desired region of interest in the brain.

The fitness or objective function used to achieve optimization is given by the ratio shown in Equation (4), which is calculated for each electrode combination and current combination:

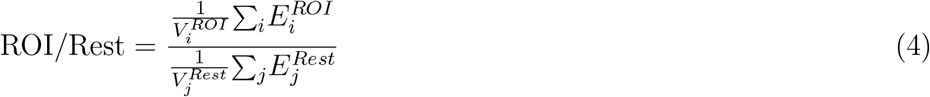

where 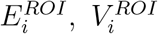 and 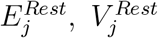 are the electric field and volume in the region of interest (ROI) and the rest of the brain areas, respectively. (A short algorithm in the supplementary material shows the calculation of the fitness function for the optimization)

### 2.5. Data Analysis

Before running the optimization, all models were numerically solved for the 61 electrodes, using the extra electrode on the left mastoid as the reference. To minimize the search space and speed up the optimization process, the currents were chosen to have a sum of 2mA, with a step of 0.05mA, with the minimum and maximum current being 0.5mA and 1.5mA, respectively [10]. The genetic algorithm was initiated with random conditions (random pairs of electrodes) and had a population of 100 chromosomes (combinations), terminating at 25 generations (iterations). The elitist version of the algorithm was used, with the elitist ratio set to 0.01 [27].

Finally, a constraint on the maximum electric field in the target region was set. Each electrode combination stimulating below the set constraint threshold was given a large penalty [27] to steer the algorithm away from the low-intensity electric field values in the target. As discussed in [10], the set constraint values do not constitute a definite threshold. The effectiveness of the optimization method is presented in the Results and Discussion section for various values (0.2, 0.5, and 0.8V/m) imposed as a maximum electric field constraint. The models in the un-optimized case were stimulated using the PO7-F7 and P8-FC6 electrode pairs, based on the 10-10 EEG system, and 1.25, 0.75mA injected current, respectively. These electrodes and current values were selected based on the H_sub2_ optimized model of Lee et al. [10].

## 3. Results and Discussion

To comparatively assess the optimization results, the left and right regions of the thalamus and hippocampus were used. Since the target region is the right hippocampus, the stimulation in all other regions is expected to be minimal, that is, ideally zero. However, zero stimulation in the regions outside the ROI cannot be achieved with this method, as the resolution provided by the electrode montages is far less than that required to achieve pinpoint accuracy.

An overview of the optimization effectiveness is presented by correlating all models in a voxel-to-voxel scheme and calculating the Pearson’s correlation coefficient, as shown in Figure 2. It can be recognized in Figure 2 that for the un-optimized case (upper left subplot) there is some variability between single subjects, as described in Conta et al. [28]. The variability in the optimized cases increased considerably since the electrode montages and current settings were not the same for all the subjects.

**Figure 2.**
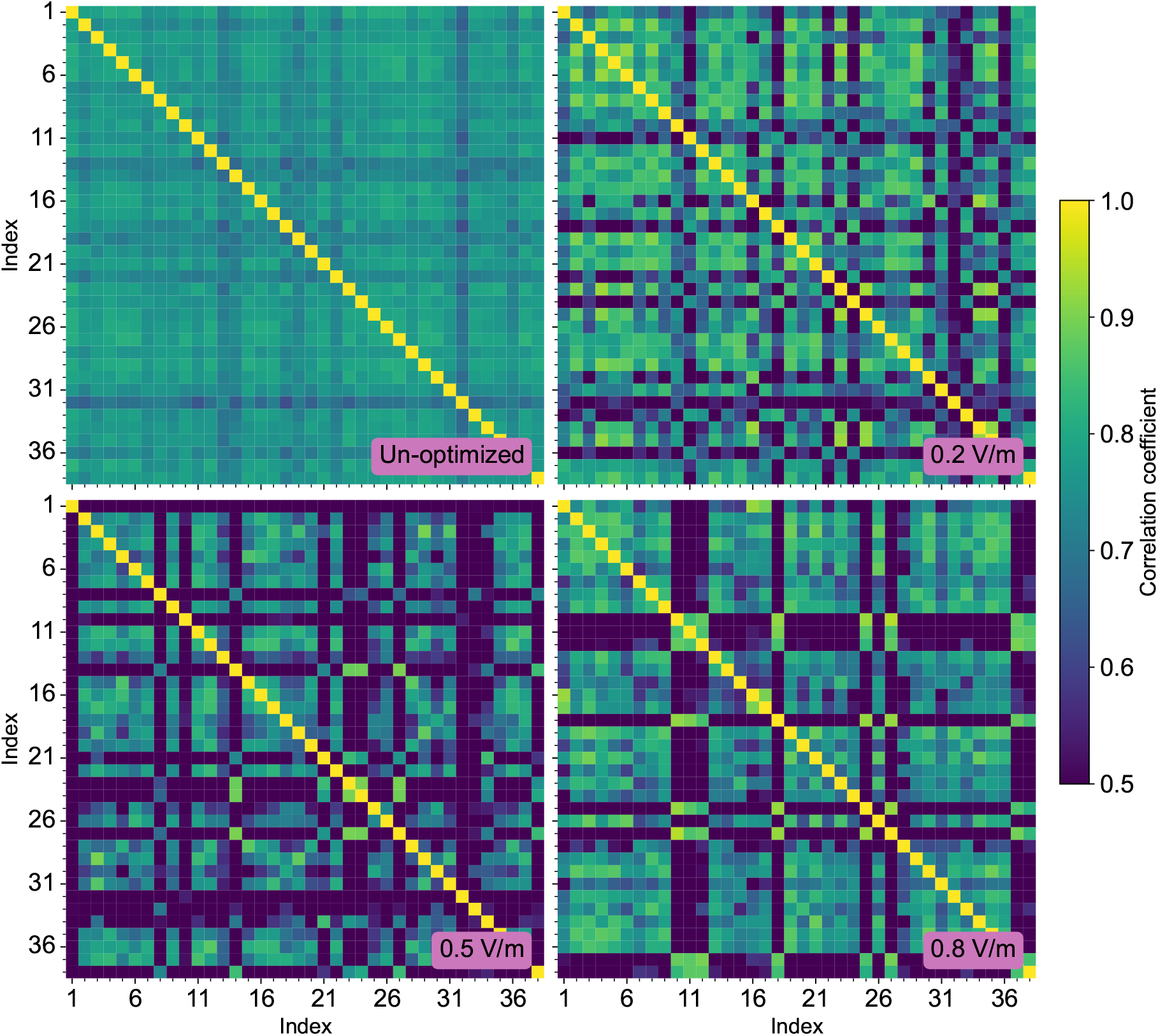
Correlation of the modulated electric field intensity across all models listed in Table 1. Only the areas identified AAL areas were considered in the correlation. The purple box at the lower right corner of each diagram indicates the optimization threshold. The colorbar shows the correlation coefficient.

All model voxels (skin, skull, CSF, etc.) other than those in the identified AAL areas were discarded, and the data calculations were performed strictly using the AAL areas. The optimization results are presented with the 0.2, 0.5, and 0.8V/m thresholds imposed for the genetic algorithm penalty function (figures 5–8).

### 3.1. Focality and Field Distribution

Establishing a focality measure will aid in assessing the quality of regional targeting in any of the cases discussed in this study. We used the modulated electric field value of 0.2V/m as the stimulation threshold, based on Lee et al. [10]. However, as Lee et al. [10] discussed, the 0.2V/m threshold is not definite as it potentially varies across individuals [29], [30].

Based on the 0.2V/m stimulation threshold, Figure 3 demonstrates that in the unoptimized case, the stimulated volume inside the ROI is comparable to that outside the ROI. In contrast, there was a significant decrease in the stimulated volume outside the ROI in the optimized cases. The optimal stimulation, considering all cases, is achieved by the 0.8V/m optimization threshold because it has the lowest spread (in the ROI) compared to all the other thresholds.

**Figure 3.**
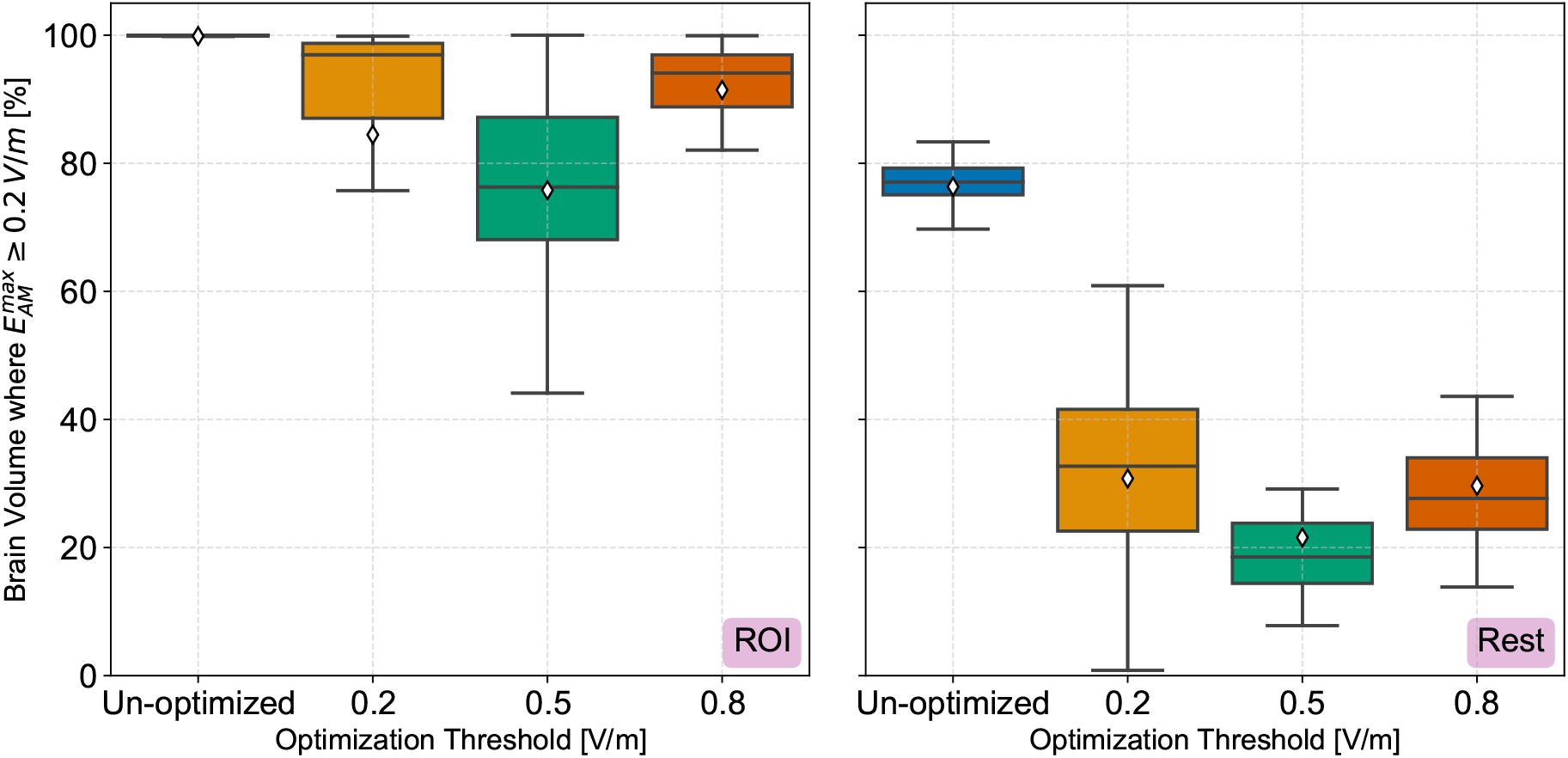
Percentage of the brain volume where 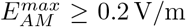 for the optimized and un-optimized stimulation of hippocampus, across all models. The subplot on the left shows the percentage inside the region of interest (hippocampus), while the subplot on the right shows the volume percentage on the rest of the regions, excluding the hippocampus. The ♢ indicator represents the arithmetic mean.

Another measure that can be used is Vol_80_, which indicates the total volume of the ROI where the condition 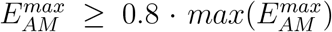 is true. Since the 0.8V/m optimization threshold is the best achieved result, we will only consider this threshold for Vol_80_ calculations. For all optimized models with the 0.8V/m as the optimization threshold, the maximum Vol_80_ volume was 427 mm^3^ (model index 24), the minimum was 35 mm^3^ (model index 5), and the mean was 190.9 mm^3^. For comparison, in the un-optimized case, the maximum Vol_80_ volume was 443 mm^3^ (model index 20), the minimum was 7 mm^3^ (model index 12), and the mean volume was 74 mm^3^.

We chose the nine best models based on the ROI/Rest ratio in descending order for the 0.8V/m optimization threshold to demonstrate the field distribution in the unoptimized and optimized cases. In Figure 4, a coronal section of the numerical difference between the un-optimized and optimized fields is presented for the nine models. As shown in Figure 4, the electric field in the optimized case focuses more on the right side (not necessarily on the ROI) than in the un-optimized case, which has a wider spread at the particular slice. While right-side focusing is the desired result of the optimization, as the target is the right hippocampus, it can be observed in Figure 4 that the electric field is not sufficiently concentrated in the ROI. Considering that we used two electrode pairs in this study, the focality resolution was limited. This problem can be addressed by using more electrode pairs for stimulation [31].

**Figure 4.**
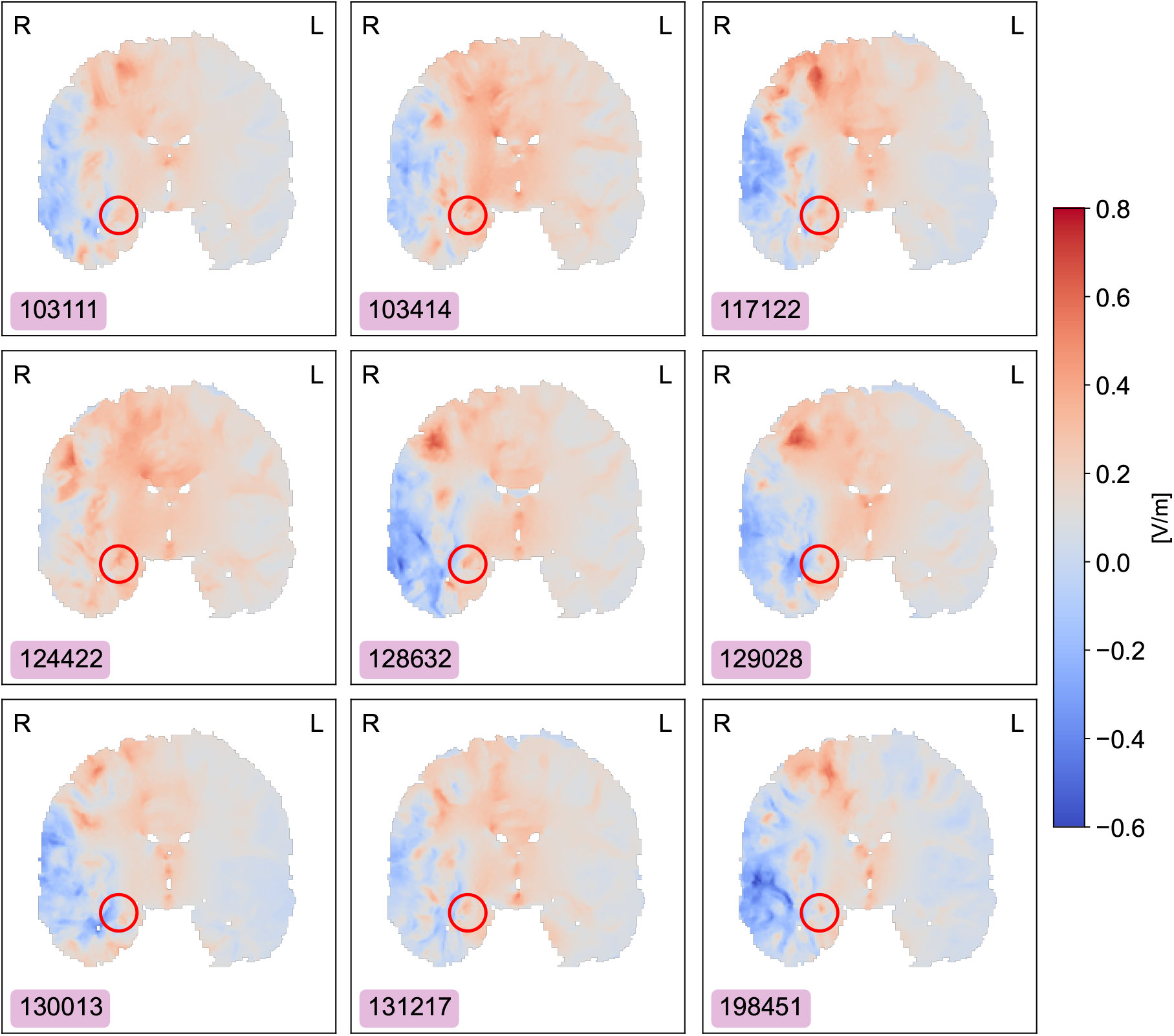
Coronal section depicting the numerical difference between the unoptimized and the optimized (for the 0.8V/m threshold) field distribution. Each model’s ID is displayed at the lower right corner and the corresponding model index can be found in Table 1. The red circle in each image indicates the area of the right hippocampus and the hemisphere is indicated at the upper right and left corners.

### 3.2. Optimization Effectiveness

The first measure of optimization effectiveness is given by Equation (4) for every optimization threshold (a constraint on the maximum modulated electric field intensity). From Figure 5, it can be observed that the ratio does not vary considerably between different thresholds, indicating that the ability to focus on the electric field is independent of the maximum value achieved.

**Figure 5.**
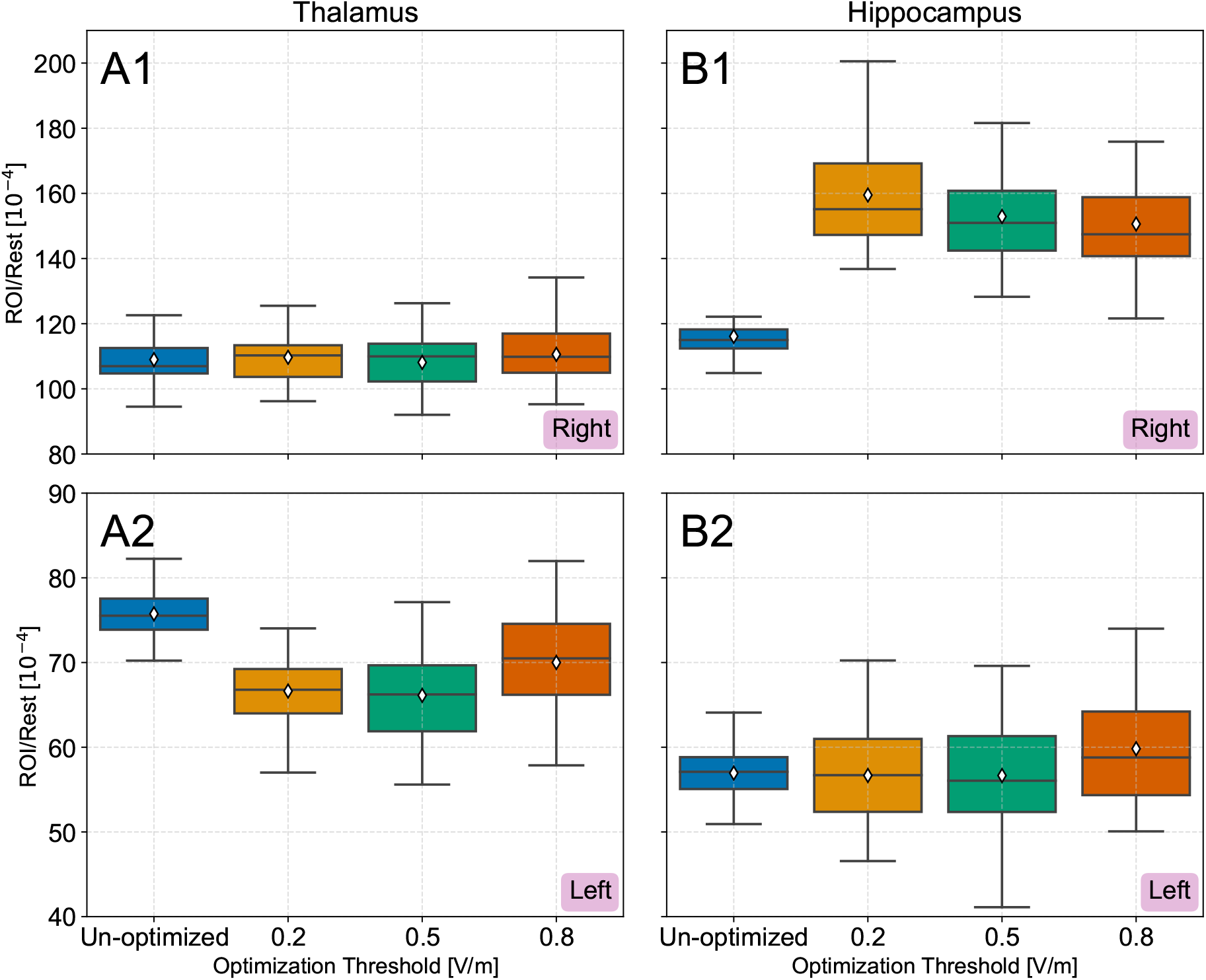
The ROI-Rest ratio for the optimized and un-optimized stimulation of the regions of thalamus and hippocampus, across all models. Subplots having the letter (A) show the values for the thalamus, while those with letter (B) show the values for the hippocampus. In all subplots the indicator at the lower right corner is the hemisphere to which the region belongs. The ♢ indicator represents the arithmetic mean.

On the other hand, Figure 6 shows that there is high variability of the maximum electric field when using the 0.2V/m threshold. This behavior is expected since the modulated electric field deep in the brain is about 0.36V/m per 1mA of injected current [8]; thus, higher optimization thresholds will provide more stable and accurate optimization results.

**Figure 6.**
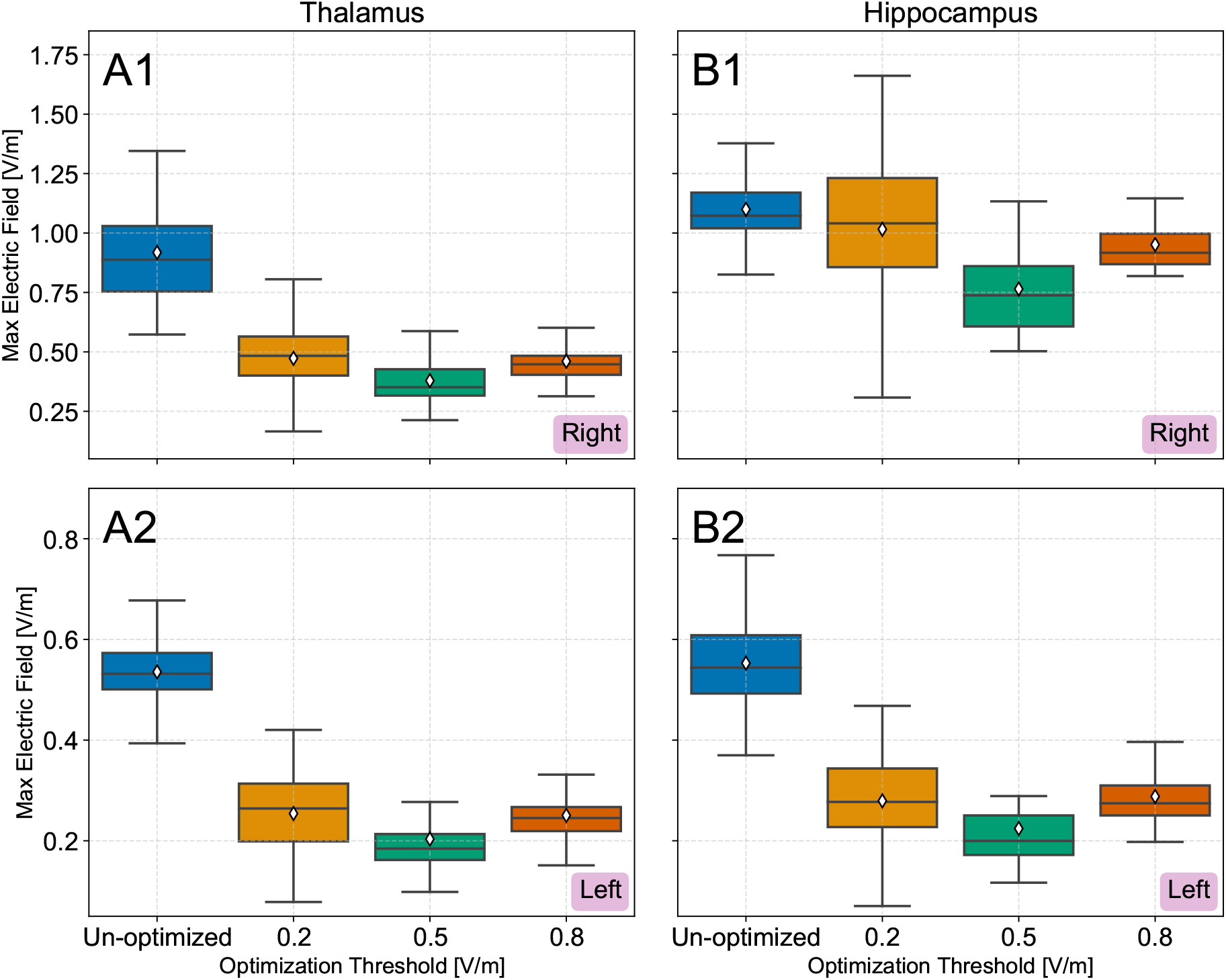
The maximum electric field value for the optimized and un-optimized stimulation of the regions of thalamus and hippocampus, across all models. Subplots having the letter (A) show the values for the thalamus, while those with letter (B) show the values for the hippocampus. In all subplots the indicator at the lower right corner is the hemisphere to which the region belongs. The ♢ indicator represents the arithmetic mean.

### 3.3. Optimization Variability

To better illustrate the optimization effect on each individual model used in this study, a comparison of the thalamus and hippocampus regions is presented in Figure 7 and 8. Each point represents one of the 38 models, with the data sorted in ascending order to make the comparison easier. Thus, for the same index, the values do not necessarily belong to the same model. (A 1-to-1 correlation of the points for the different optimization thresholds can be found in the supplementary material.)

**Figure 7.**
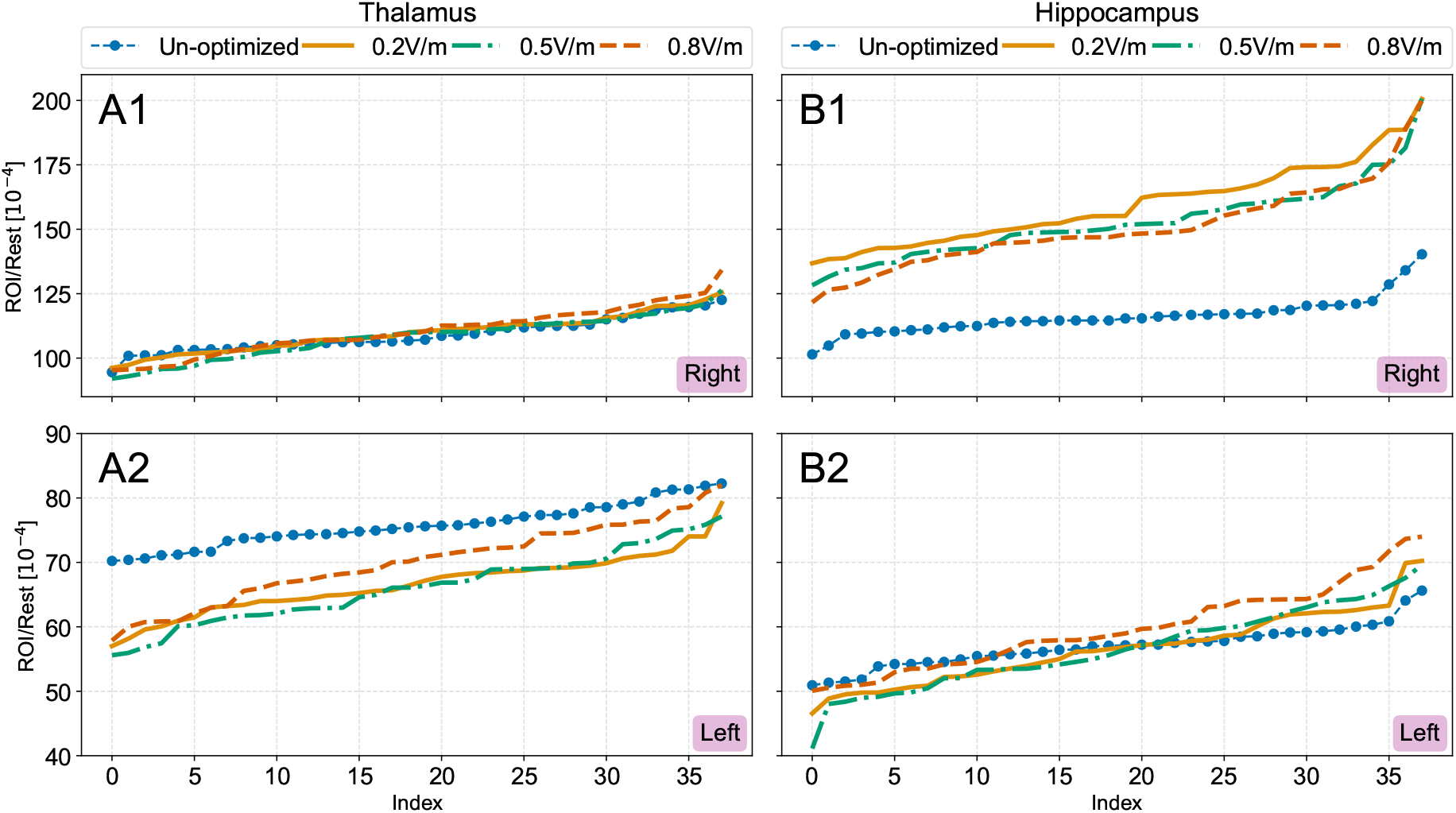
The ROI-Rest ratio for the optimized and un-optimized stimulation of the regions of thalamus and hippocampus, for each individual model. Subplots having the letter (A) show the values for the thalamus, while those with letter (B) show the values for the hippocampus. In all subplots the indicator at the lower right corner is the hemisphere to which the region belongs. The data points for each subplot are sorted for visualization purposes, meaning that the data represented by the same index are not necessarily correlated. The index on the x axis is the ascending count of the models, with no relevance to Table 1.

**Figure 8.**
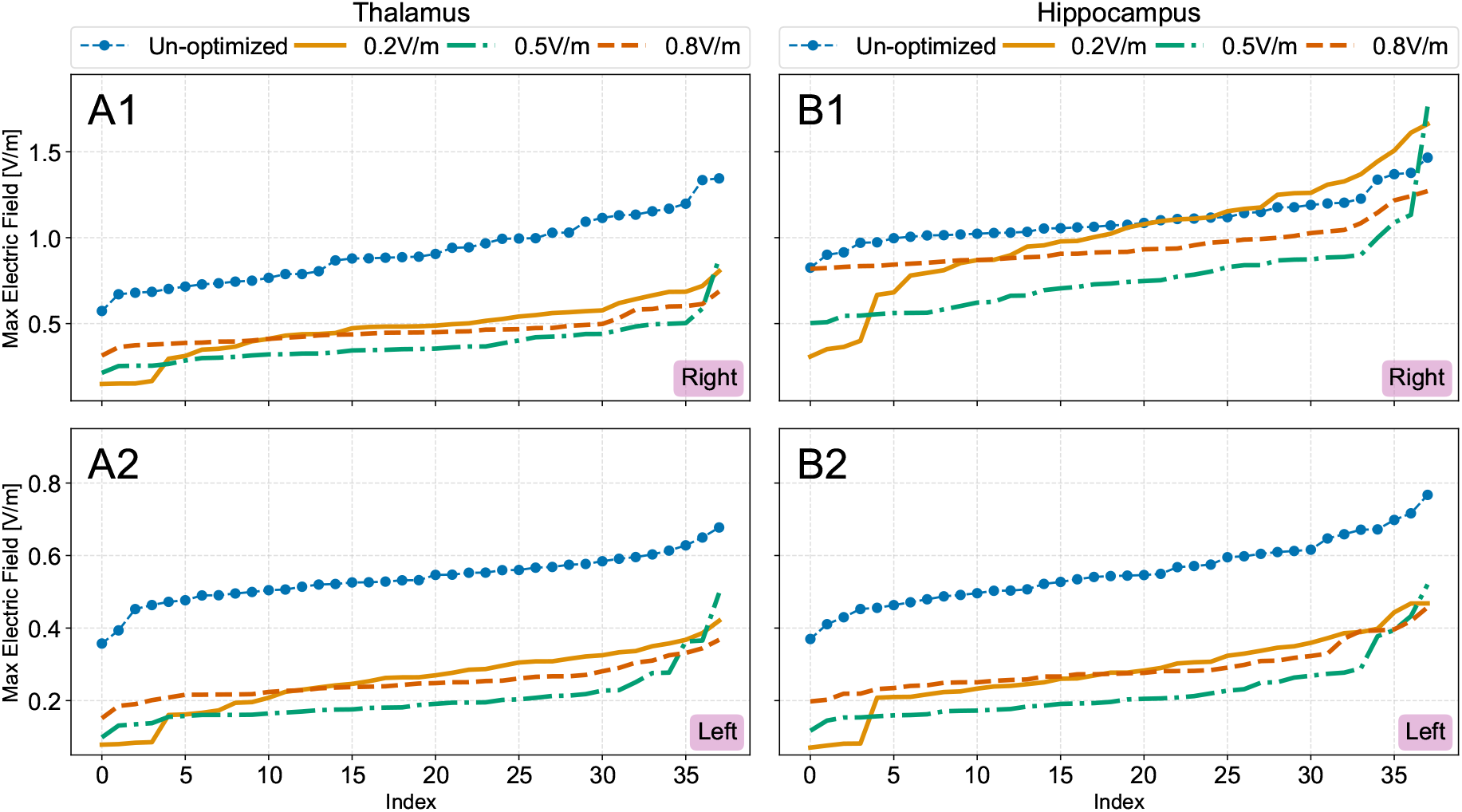
The maximum electric field value for the optimized and un-optimized stimulation of the regions of thalamus and hippocampus, for each individual model. Subplots having the letter (A) show the values for the thalamus, while those with letter (B) show the values for the hippocampus. In all subplots the indicator at the lower right corner is the hemisphere to which the region belongs. The data points for each subplot are sorted for visualization purposes, meaning that the data represented by the same index are not necessarily correlated. The index on the x axis is the ascending count of the models, with no relevance to Table 1.

As shown in Figure 7, using Equation (4) as the optimization metric across all models, there is a minimal change in the different thresholds. The right hippocampus in all optimized cases had a significantly higher ratio than the left hippocampus, indicating a stronger field concentration in the right hippocampal area, which is also supported by Figure 3.

Same as in Figure 6, it can be observed in Figure 8 that, as the threshold of the penalty function increases, the spread of the maximum value of the modulated electric field decreases. The wide variations in the right hippocampus for the 0.2V/m threshold, can be attributed to the low value of the threshold, as the optimization algorithm is unable to successfully meet the optimization criteria. Since the total injected current from both electrode pairs is 2mA, following the 0.36V/m per 1mA rule [8], the expected electric field values are larger than the constraint itself for the 0.2 and 0.5V/m thresholds; thus, the algorithm may never approach an optimal solution.

Optimization profoundly affects the maximum electric field value and distribution across the region of interest. The higher the optimization threshold, the smaller the spread of the maximum electric field observed between the models, leading to a more focused stimulation than the un-optimized stimulation.

Figure 8 indicates that stimulation in the region of the right thalamus does not differ much from the stimulation in the region of the right hippocampus, although the latter has a higher maximum electric field value. This result is explained by the size difference between the thalamus and hippocampus (the thalamus is larger than the hippocampus) and the relative position of the structures (structures are close to each other).

Given that the accomplishment of a globally optimal solution consistently requires a brute-force approach, utilizing the genetic algorithm can significantly decrease the optimization time and provide meaningful results in less than two hours for each model.

### 3.4. Method Improvements and Considerations

There have been studies regarding the robustness and effectiveness of the tTIS method, trying to resolve focality issues. A multi-channel tTIS study on mice [31] indicated that targeting accuracy significantly improves when considering a six-channel (six electrode pairs) tTIS, performing 70.2% better than the single-channel (one electrode pair) tTIS. Furthermore, in [31], stimulation targeting appeared to be less affected when selecting different electrode pairs. In summary, Song et al. [31] showed that more channels make the tTIS method more independent of the exact electrode placement while increasing the focality. Studying the multi-channel approach on human head models and ultimately conducting clinical trials will shed light on potential improvements.

Additionally, as discussed and illustrated in [9], there is strong evidence [32], [33] that neurons respond preferentially to stimulation and that preference seems to be oriented along the neuron’s axon direction. Comparing the field distribution in Figure 4 with figures 5 and 11 from [9], it is evident that in the case where the neuron stimulation preference is not considered, the field has a more extensive spread within the brain. Combining the neurons’ preferential nature and the multi-channel approach, there are strong indications that the results will be significantly better at the individual level.

## 4. Conclusion

In this study, we applied the temporal interference stimulation (tTIS) method, using two electrode pairs, on 38 numerical head models of the PHM repository. Each model was fitted with electrodes according to the 10-10 international EEG electrode placement system. We optimized the electrode combinations and the injected current on each pair and showed that it is possible to achieve targeted stimulation on individual models. More specifically, from our analysis, it is evident that greater focality is achieved, considering the right hippocampus as the region of interest, with a significant decrease in the surrounding electric field intensity.

In addition, a diminishing spread of the maximum modulated electric field values in the optimized models was observed as the optimization threshold increased. In contrast, in the un-optimized case (where the same electrode pairs and currents are used for all models), the modulated electric field across the models has relatively low variability, and region targeting is not optimal. Additionally, we observed that the selected objective function value for the optimization runs seems to be independent of the maximum electric field value achieved in the target region. In some cases, the electric field outside the region of interest is comparable to that inside the ROI, suggesting that achieving focused stimulation in all subjects with the tTIS method is difficult, mainly because of the small target volume.

Apart from purely anatomical constraints, using two electrode pairs is also a limiting factor since the electric field outside the target region is not easily steerable. Potentially utilizing more electrode pairs can solve the focality issues. Finally, the results from the genetic algorithm in this study are promising, indicating that individual optimization can be achieved without using exhaustive search methods.

## 6. Data and Software Availability

The software used in this publication are available on Zenodo [34].

## Acknowledgments

Results presented in this work have been produced using the Aristotle University of Thessaloniki (AUTh) High Performance Computing Infrastructure (https://it.auth.gr/en/hpc).

