## Supplementary Material (PDF) for "Non-invasive stimulation with Temporal Interference: Optimization of the electric field deep in the brain with the use of a genetic algorithm"

Theodoros Samaras

### Contents

**1 Material Description** **2**

**2 Electrode Positions and Currents** **3**

**3 Scatter Plots Across all Models** **6**

**4 Optimization Objective Function Algorithm** **7**

**References** **8**

### 1 Material Description

In section 2, all the electrode combinations and currents for all three optimization thresholds are presented in figures S3 - S5. Each electrode pair is given a designator, so two electrodes with the same designator correspond to the same pair. The designators are △ and □. The color of each of the designators in the figures corresponds to the same colored current value on the lower left and right corners, with the ground electrode colored black. Moreover, the electrodes and currents used in the un-optimized case are depicted in figure S1b. All the electrode names at each position can be found in figure S1a.

Figure S2 compares our results (subfigure S1a) with those of Grossman et al. [1, Figure S2Ki] (subfigure S1b). The electrodes were placed according to figure 2C of Grossman et al. [1] and the stimulation current was 1 mA at each electrode pair. (Code available on Zenodo [2])

Section 3 shows the scatter plots for the ROI/Rest and maximum electric field values for all cases. Lastly, in section, 4 the algorithm used to calculate the optimization objective function can be found.

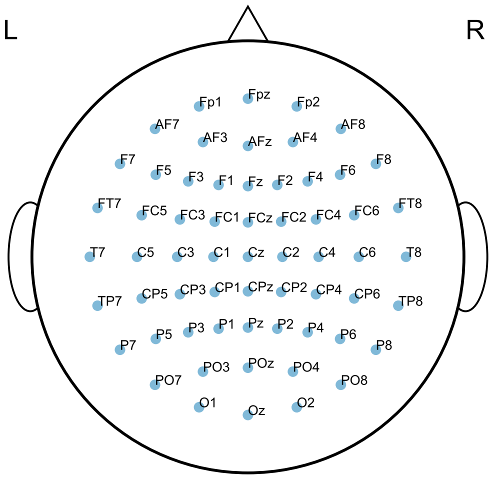


**(a)** Electrode names of the 10-10 EEG electrode placement system.

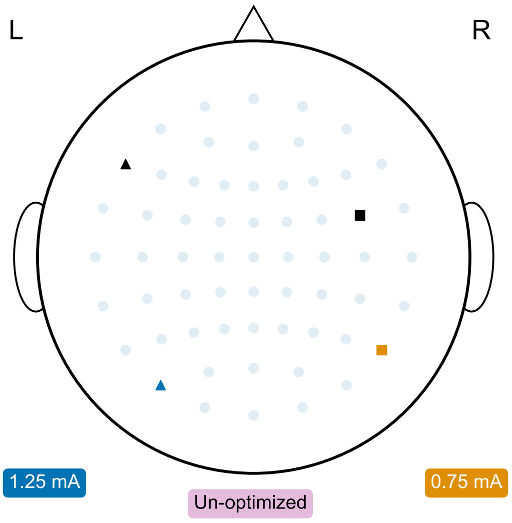


**(b)** Electrode locations and currents for the un-optimized case.

**Figure S1:** Electrode names (a), electrode and current combination for the un-optimize case (b).

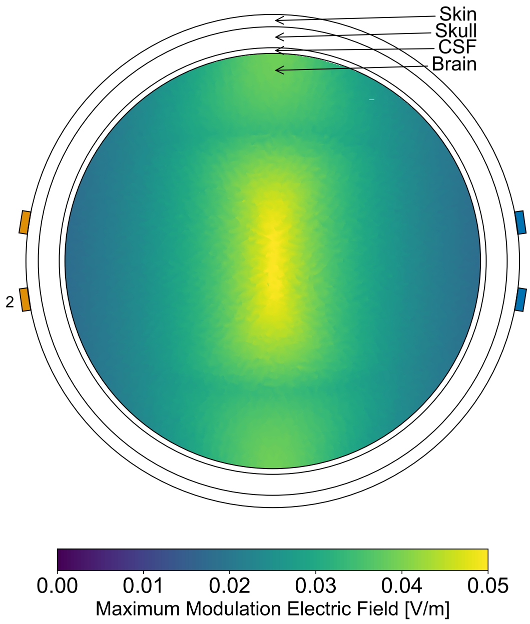


**(a)** Our software.

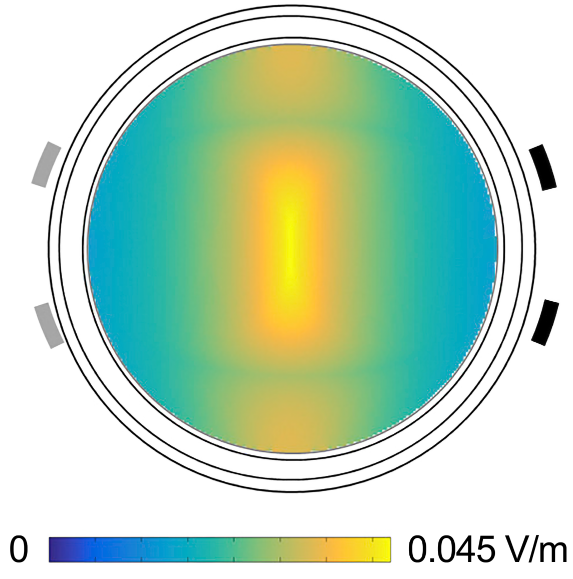


**(b)** Figure S2Ki of Grossman et al. [1].

**Figure S2:** Software results comparison. Is subfigure (a) the results from our work are presented, while in subfigure (b) the reusults of Grossman et al. [1, Figure S2Ki] are displayed for comparison.

### 2 Electrode Positions and Currents

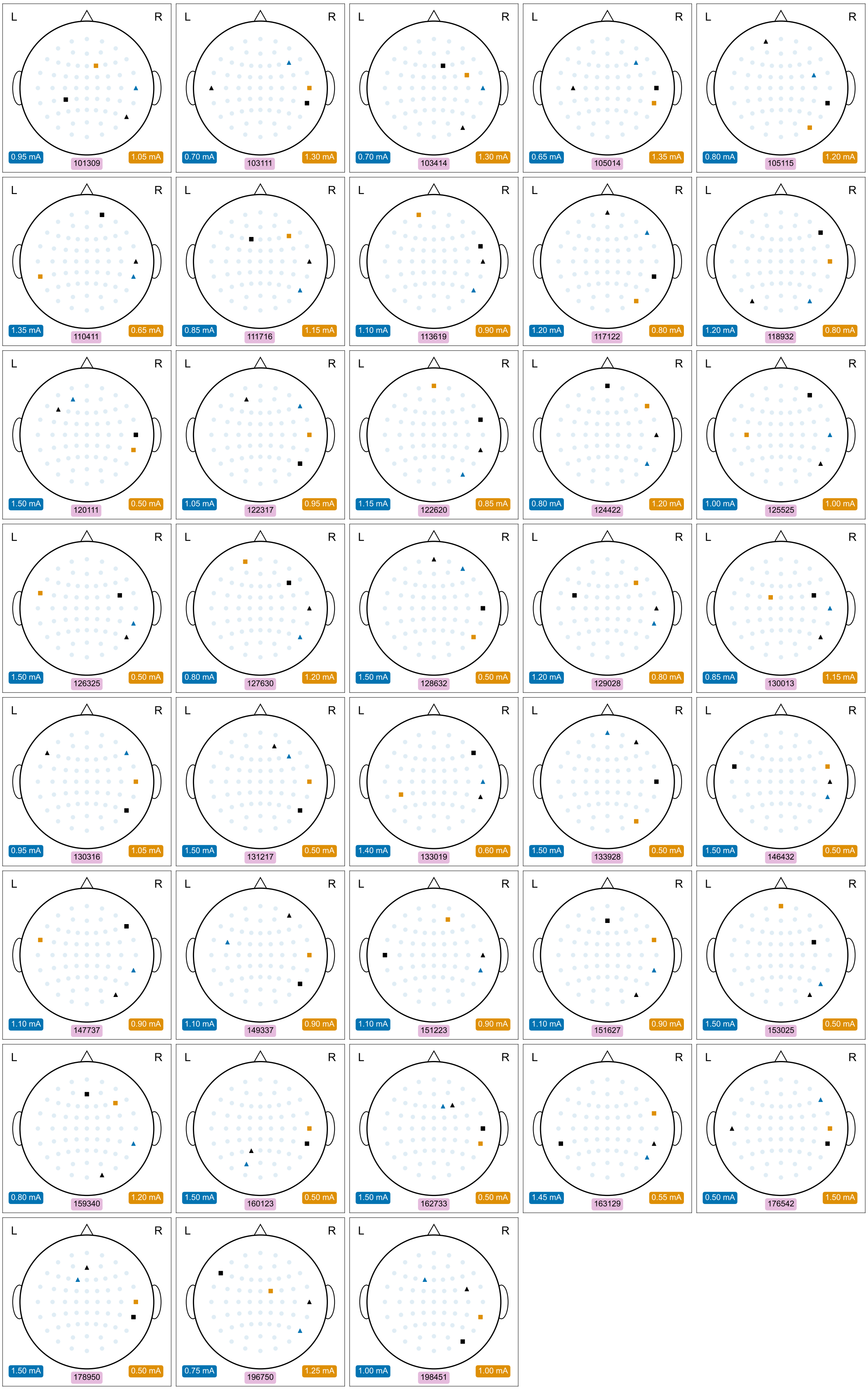


**Figure S3:** Optimized electrode positions for the 0.2 V/m threshold. Currents shown on the lower left corner correspond to the △ designator pair, while those on the lower right correspond to the □. The black colored designators indicate the ground. Additionally, the model ID is displayed in the middle.

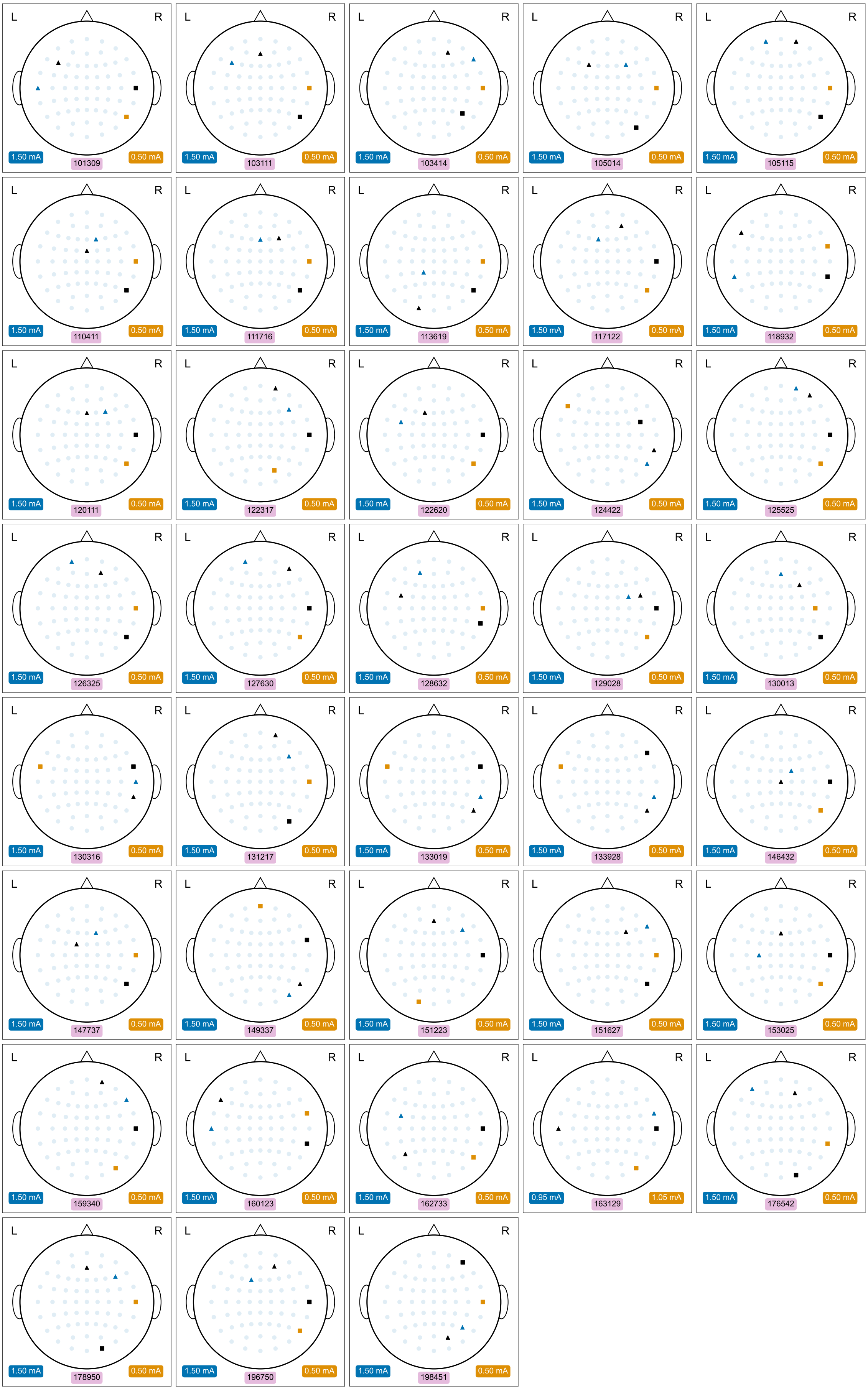


**Figure S4:** Optimized electrode positions for the 0.5 V/m threshold. Currents shown on the lower left corner correspond to the △ designator pair, while those on the lower right correspond to the □. The black colored designators indicate the ground. Additionally, the model ID is displayed in the middle.

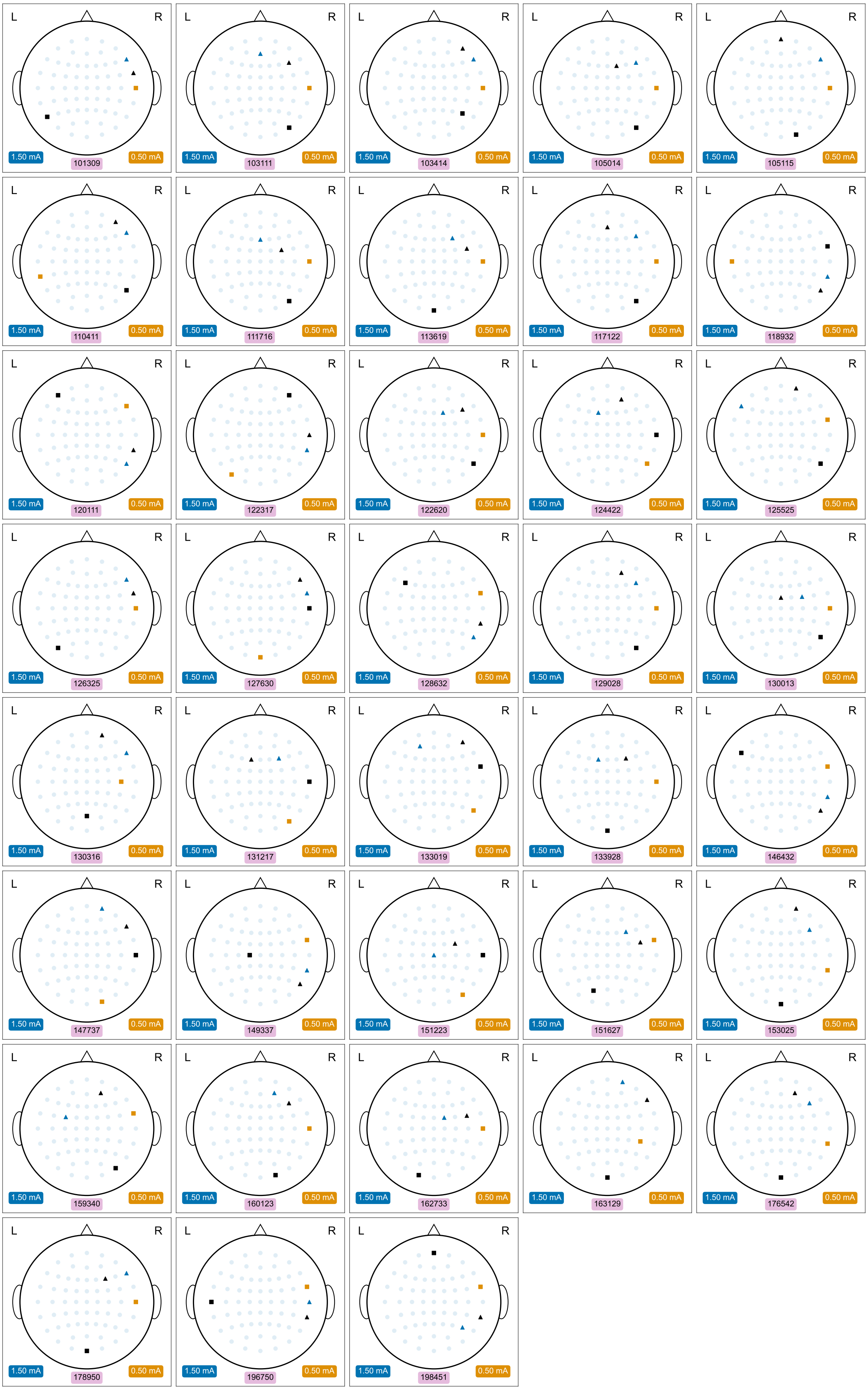


**Figure S5:** Optimized electrode positions for the 0.8 V/m threshold. Currents shown on the lower left corner correspond to the △ designator pair, while those on the lower right correspond to the □. The black colored designators indicate the ground. Additionally, the model ID is displayed in the middle.

### 3 Scatter Plots Across all Models

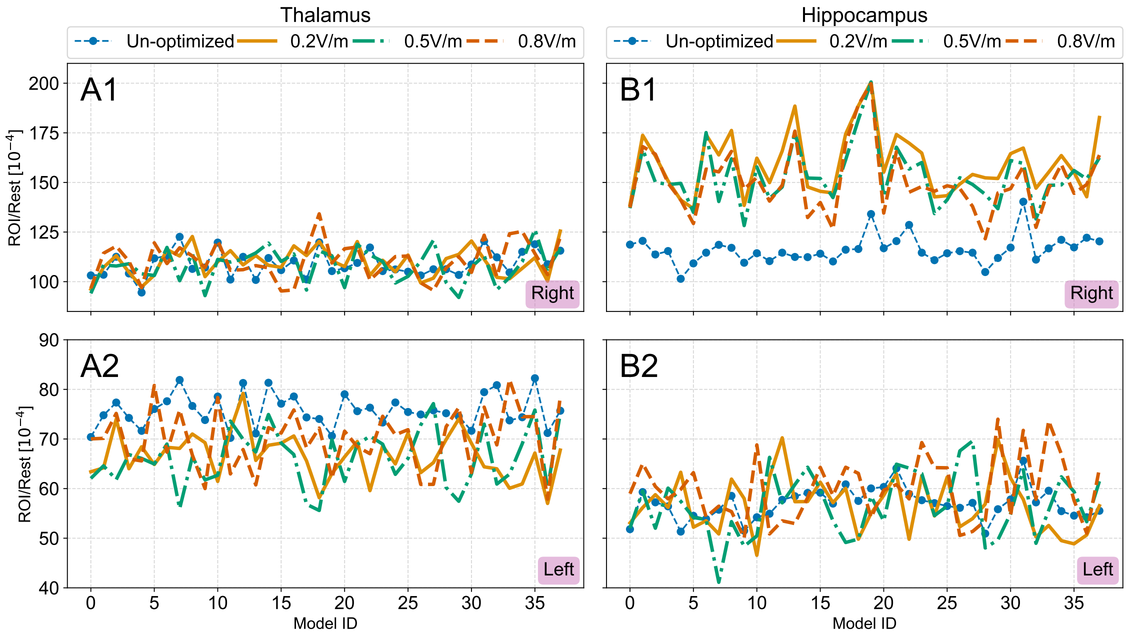


**Figure S6:** The ROI-Rest ratio for the optimized and un-optimized stimulation of the regions of thalamus and hippocampus, for each individual model. Subplots having the letter (A) show the values for the thalamus, while those with letter (B) show the values for the hippocampus. In all subplots the indicator at the lower right corner is the hemisphere to which the region belongs. The data points for each subplot are 1-1 correlated.

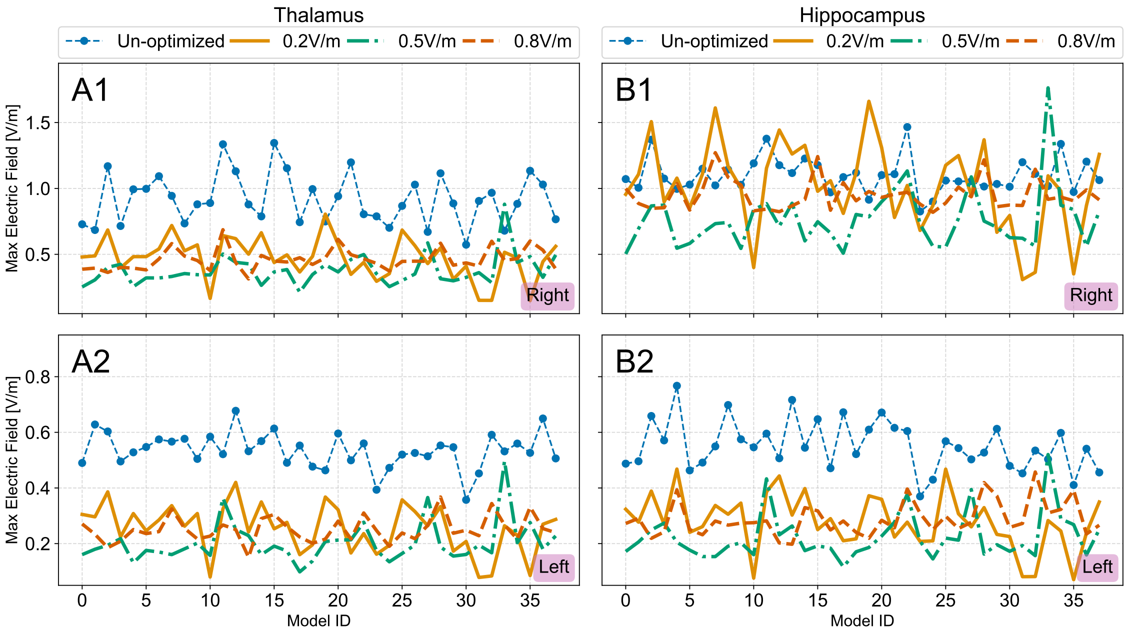


**Figure S7:** The maximum electric field value for the optimized and un-optimized stimulation of the regions of thalamus and hippocampus, for each individual model. Subplots having the letter (A) show the values for the thalamus, while those with letter (B) show the values for the hippocampus. In all subplots the indicator at the lower right corner is the hemisphere to which the region belongs. The data points for each subplot are 1-1 correlated.

### 4 Optimization Objective Function Algorithm

**Algorithm 1:** Optimization objective function value calculation steps.

**Input :** A list of integers $[e_i]$ holding the index of each electrode, an array $[field\_data_{ij}]$ containing the solved electrodes based on the ground reference, an array $[regions\_of\_interest_i]$ containing the regions of interest for the optimization, an array $[aal\_regions_i]$ containing the corresponding AAL atlas region for each electric field point, a dictionary $[region\_volumes_i]$ containing the total volume of each AAL region and an array $[currents_{ij}]$ containing all the valid current combinations. Lastly, the desired optimization threshold $opt\_thr$ is give.

**Output:** The fitness value of the optimization.

**if** *All* $[e_i]$ *integers are not unique* **then**
    **return** $100 * abs(length(unique([e_i])) - 4)^2 + 10000$;
**end if**
$penalty \longleftarrow 0$;
$roi \longleftarrow$ Set to $True$ if $[aal\_regions_i]$ and $[regions\_of\_interest_i]$ match, otherwise set to $False$;
Set an empty list $[max\_vals_i]$ and $[fitness\_vals_i]$ to hold the maximum modulation values and the fitness values, respectively;
**foreach** $current \in [currents_i]$ **do**
    $e\_field\_base \longleftarrow current[0] * field\_data[e[0]] - current[0] * field\_data[e[1]]$;
    $e\_field\_df \longleftarrow current[1] * field\_data[e[2]] - current[1] * field\_data[e[3]]$;
    $modulation\_values \longleftarrow$ modulation_envelope($e\_field\_base$, $e\_field\_df$);
    Append to $[max\_vals_i]$ the max($modulation\_values[roi]$);
    $roi\_region\_sum \longleftarrow 0$;
    $non\_roi\_region\_sum \longleftarrow 0$;
    **foreach** $region \in unique([aal\_regions_i])$ **do**
        $roi \longleftarrow$ Find where $[aal\_regions_i] == region$, for every $i = 1, 2, \cdots, n$;
        **if** $region \in [regions\_of\_interest_i]$ **then**
            $roi\_region\_sum \longleftarrow roi\_region\_sum + sum(\text{modulation\_vlues}[roi])/region\_volumes[region]$;
        **else**
            $non\_roi\_region\_sum \longleftarrow roi\_region\_sum + sum(\text{modulation\_vlues}[roi])/region\_volumes[region]$;
        **end if**
    **end foreach**
    $region\_ratio \longleftarrow roi\_region\_sum/non\_roi\_region\_sum$;
    $fitness\_measure \longleftarrow region\_ratio * 10000$;
**end foreach**
Append to $[fitness\_vals_i]$ the $fitness\_measure$;
$max\_val\_curr \longleftarrow$ Set to $True$ where $[max\_vals_i] >= opt\_thr$, for every $i = 1, 2, \cdots, n$, otherwise set to $False$;
$return\_fitness \longleftarrow 0$;
**if** *max_val_curr does not have True values* **then**
    $penalty \longleftarrow penalty + 100 * (opt\_thr - mean(max\_vals))^2 + 1000$;
    $return\_fitness \longleftarrow min(fitness\_vals)$;
**else**
    $return\_fitness \longleftarrow max(fitness\_vals[max\_val\_curr])$;
**end if**
**return** $-(return\_fintness - penalty)$
