## Supplementary figures and images for "Non-invasive stimulation with Temporal Interference: Optimization of the electric field deep in the brain with the use of a genetic algorithm"

### Supplemental Figure S3

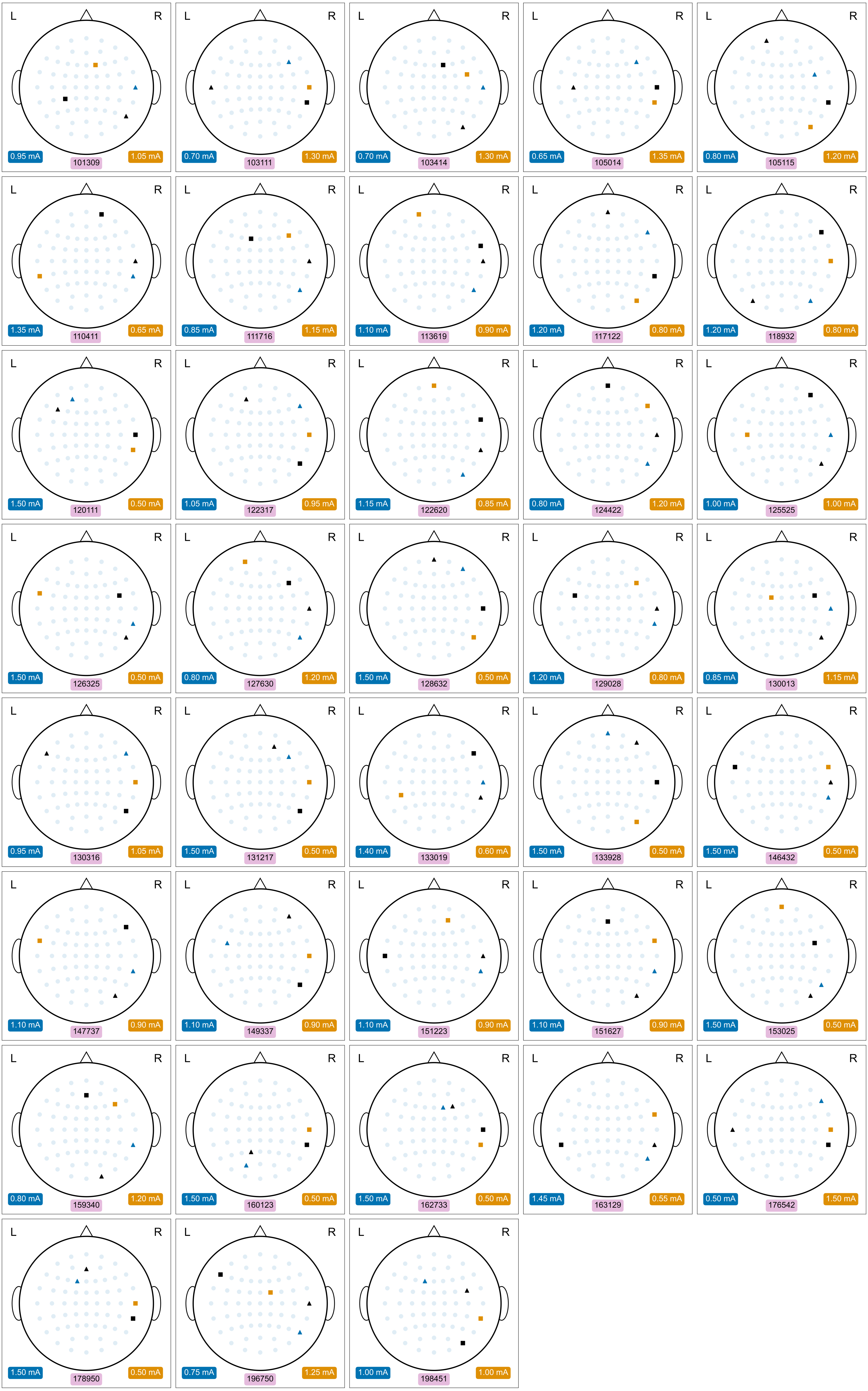

### Supplemental Figure S4

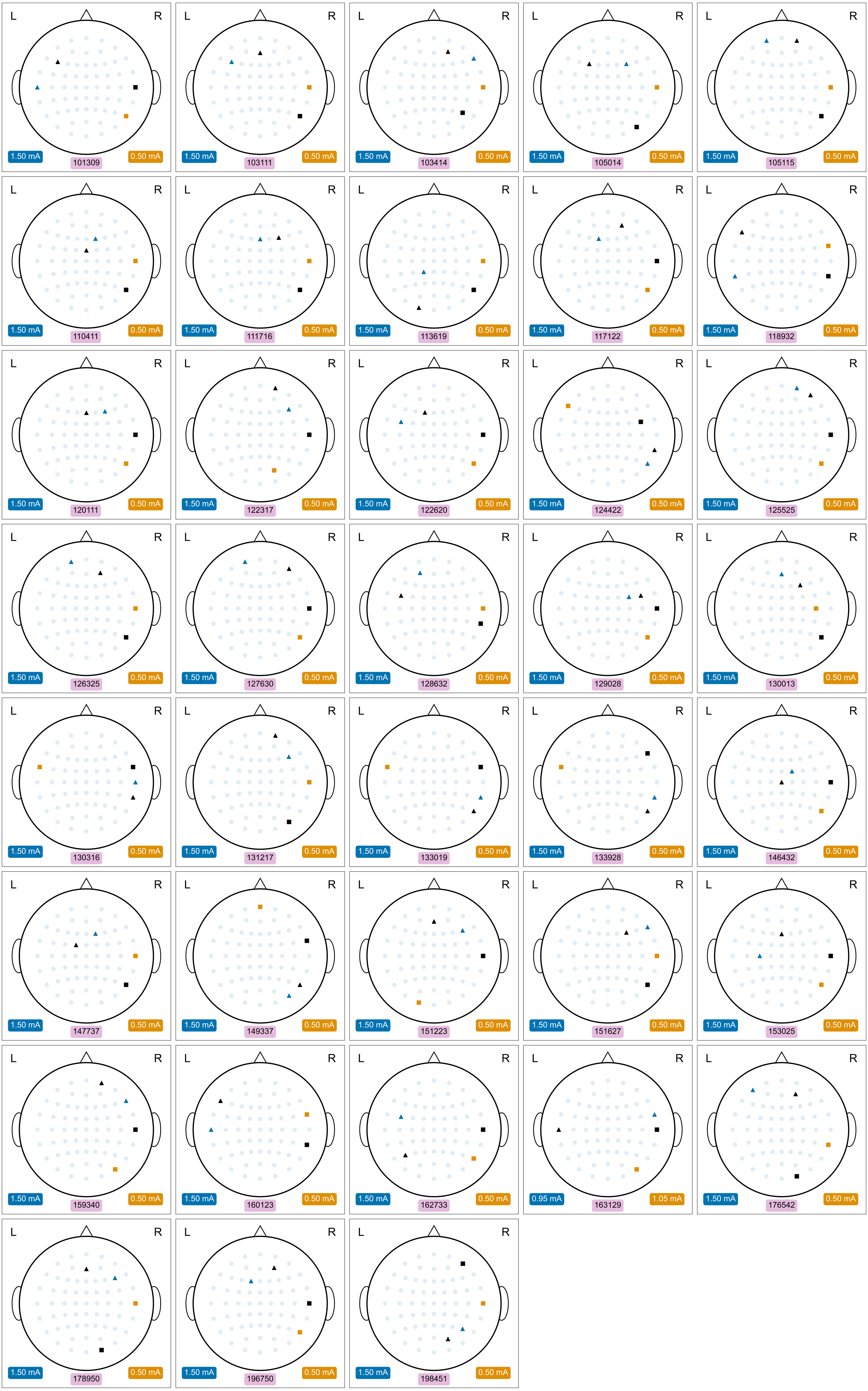

### Supplemental Figure S5

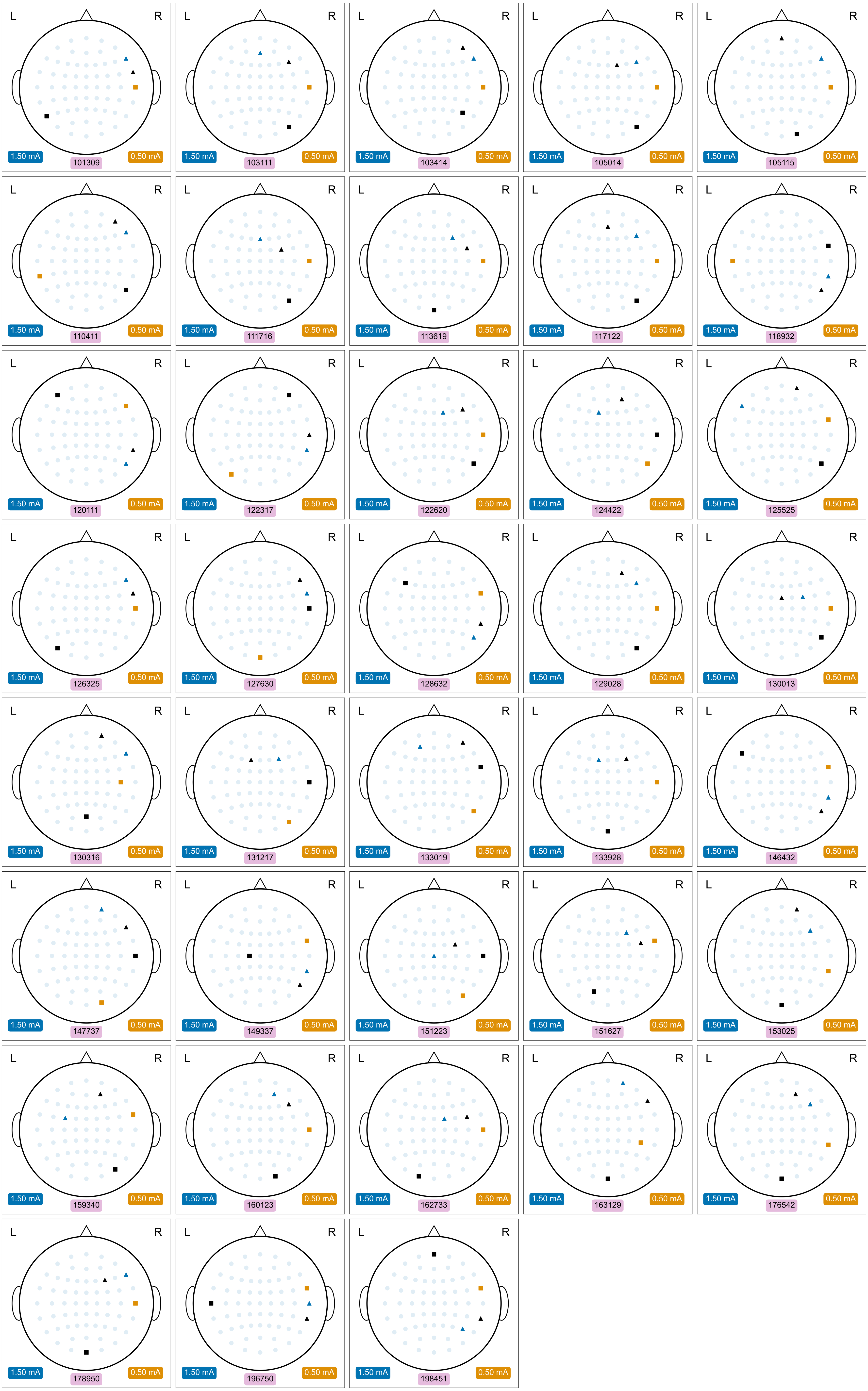
